# The complete genome sequence of the crayfish pathogen Candidatus *Paracoxiella cheracis* n.g. n.sp. provides insight into pathogenesis and the phylogeny of the Coxiellaceae family

**DOI:** 10.1101/2024.12.02.626477

**Authors:** Danielle J. Ingle, Calum J. Walsh, Genevieve R. Samuel, Ryan R. Wick, Nadav Davidovich, Eleonora Fiocchi, Louise M. Judd, Jennifer Elliman, Leigh Owens, Timothy P. Stinear, Andrea Basso, Tobia Pretto, Hayley J. Newton

## Abstract

The Coxiellaceae bacterial family, within the order Legionellales, is defined by a collection of poorly characterized obligate intracellular bacteria. The zoonotic pathogen and causative agent of human Q fever, *Coxiella burnetii*, represents the best characterized member of this family. Coxiellaceae establish replicative niches within diverse host cells and rely on their host for survival, making them challenging to isolate and cultivate within a laboratory setting. Here we describe a new genus within the Coxiellaceae family that has been previously shown to infect economically significant freshwater crayfish. Using culture-independent long-read metagenomics, we reconstructed the complete genome of this novel organism and demonstrate that the previously referred to as Candidatus *Coxiella cheraxi* represents a novel genus within this family, herein denoted Candidatus *Paracoxiella cheracis*. Interestingly, we demonstrate that Candidatus *P. cheracis* encodes for a complete, putatively functional Dot/Icm type 4 secretion system that likely mediates the intracellular success of this pathogen. In silico analysis defined a unique repertoire of Dot/Icm effector proteins and highlighted homologues of several important *C. burnetii* effectors including a homologue of CpeB that was demonstrated to be a Dot/Icm substrate in *C. burnetii*.

**IMPORTANCE:** Using long-read sequencing technology we have uncovered the full genome sequence of Candidatus *Paracoxiella* cheracis, a pathogen of economic importance in aquaculture. Analysis of this sequence has revealed new insight into this novel member of the Coxiellaceae family, demonstrating that it represents a new genus within this poorly characterized family of intracellular organisms. Importantly, the genome sequence reveals invaluable information that will support diagnostics and potentially both preventative and treatment strategies within crayfish breeding facilities. Candidatus *P. cheracis* also represents a new member of Dot/Icm pathogens that rely on this system to establish an intracellular niche. Candidatus *P. cheracis* possesses a unique cohort of putative Dot/Icm substrates that constitute a collection of new eukaryotic cell biology manipulating effector proteins.

## INTRODUCTION

Australian native freshwater redclaw crayfish, *Cherax quadricarinatus*, are bred for commercial market. Aquaculture of these animals has led to the observation of infectious diseases that impact both wild and farmed *C. quadricarinatus* (reviewed in (1)). In particular, a Rickettsial-like intracellular bacterium, designated TO-98, isolated from infected crayfish was recognized as the likely causative agent of mass mortality in farmed redclaw crayfish in the early 1990s (2, 3). Subsequent investigations were able to recapitulate disease in experimentally infected crayfish and demonstrate that TO-98 causes a lethal infection in redclaw crayfish (4). This study conducted 16s rRNA sequence analysis and revealed that TO-98 was closely related to the zoonotic human pathogen *Coxiella burnetii*, coining the name Candidatus *Coxiella cheraxi* (4).

The initial observation that this pathogen is a member of the Coxiellaceae family has been confirmed by examination of additional DNA sequences including comparison of specific gene sequences (5) and MinION sequencing data showing fragments of DNA with closest similarity to *C. burnetii* and *Coxiella*-like endosymbionts (C-LEs) (6). However, this fragmented genomic data does not allow complete and comprehensive understanding of the relationship between *C. burnetii* and Candidatus *C. cheraxi*.

*C. burnetii*, the causative agent of human Q fever, has a large reservoir in domesticated ruminants (7). *C. burnetii* is a Gammaproteobacteria within the order Legionellales and family Coxiellaceae. While the order Legionellales is best known for the two major human pathogens, *Legionella pneumophila* and *C. burnetii*, these bacteria are distantly related and the order is poorly characterized, including over 450 uncultured genera (8). The diversity of Legionellales has been revealed by increased sampling of different environments and ecological surveys employing metagenomic techniques that have demonstrated the presence of Legionellales within arthropods, amoeba and aquatic environments (reviewed in (9)). Most of these newly discovered Legionellales are likely nonpathogenic bacteria that have evolved mutualistic relationships with non-vertebrate hosts. However, recent data suggests that some poorly characterized Legionellales harbor the potential to be zoonotic pathogens and/or pathogenic to important animal species. This includes a recently identified *Coxiella* species found in the placenta of fur seals (10).

Essential to the ability of *C. burnetii* and *L. pneumophila* to replicate intracellularly, and cause disease, is the Dot/Icm type IVB secretion system (T4SS) (11–14). This multiprotein apparatus is responsible for the delivery of a large cohort of effector proteins into the host cell where they modulate a range of cellular functions and pathways (reviewed in (15)). Dot/Icm-deficient *L. pneumophila* are rapidly destroyed via lysosomal degradation as many of the effectors act to remodel the phagosome and block endocytic maturation (reviewed in (16)). In contrast, internalized *C. burnetii* is trafficked through the endocytic pathway before the lysosomal environment triggers activation of the Dot/Icm system (17, 18). Dot/Icm-deficient *C. burnetii* are not destroyed by the lysosome but they are also incapable of intracellular replication (11, 12). Interestingly, comparative genomic studies show that the Dot/Icm system is lost or pseudogenized in C-LE, indicating its association with pathogenesis (19).

Despite *L. pneumophila* and *C. burnetii* relying on the same key virulence factor, the cohort of effector proteins translocated by this secretion system are pathogen specific. This genetic divergence is represented by phenotypic divergence through the establishment of distinct intracellular niches. The biochemical and functional characterization of these effector proteins is an active area of research as these novel proteins facilitate insight into both mechanisms of pathogenesis and novel strategies to manipulate eukaryotic cell biology.

Here we report the complete genome sequence of Candidatus *C. cheraxi* revealing a more distant relationship to *C. burnetii* than expected, notably that the Candidatus *C. cheraxi* represents a novel genus within Coxiellaceae. We propose to rename this crayfish pathogen Candidatus *Paracoxiella cheracis.* Interestingly, we report that this novel Coxiellaceae family member possesses a Dot/Icm T4SS most closely associated with *C. burnetii*. *In silico* analysis allowed us to identify a large cohort of putative effectors of this secretion system with only 12 out of 238 predicted effectors sharing identity to known *C. burnetii* T4SS effectors. We have been able to demonstrate translocation of the Candidatus *P. cheracis* CpeB homologue via the *C. burnetii* Dot/Icm system. This study provides additional insight into the poorly characterized Coxiellaceae family and demonstrates that the Dot/Icm secretion system is a common tool used by these bacteria to communicate with their host. It also demonstrates the utility of long-read sequencing approaches for generating a complete metagenome-assembled genome for an obligate intracellular pathogen.

## METHODS

### Collection and preparation of genetic material

Samples of hepatopancreas from infected redclaw crayfish were collected during an outbreak of crayfish rickettsiosis, which occurred after the crayfish were imported from Australia to Israel in 2019 (20). Briefly, 10,000 juvenile Redclaw crayfish (2–3 weeks old) were imported to Israel and quarantined in a facility. From April to July, juvenile mortality sharply increased despite the absence of clinical signs or gross lesions. Specimens collected for histological and molecular evaluations were sent to the Istituto Zooprofilattico Sperimentale delle Venezie in Legnaro, Italy, and revealed a severe infection by Candidatus *C. cheraxi.* DNA was extracted from these samples, as previously described (20), and sent to Monash University, Australia, for further analysis.

### Long-read sequencing and genome assembly

Metagenomic sequencing of extracted DNA was performed on two separate sequencing runs. The first run was performed on an Oxford Nanopore Technologies (ONT) MinION while the second employed a PromethION. Both runs were carried out on R10.4.1 flow cells. Basecalling and quality filtering was performed by Dorado (v. 0.5.0) using models dna_r10.4.1_e8.2_400bps_sup@v4.1.0 and dna_r10.4.1_e8.2_400bps_sup@v4.3.0 respectively. Reads shorter than 1kbp were discarded. Kraken2 (21) was used for initial read-level classification of metagenomic data.

Initial assemblies were generated using Flye (22), MetaMDBG (23), Miniasm (24), and Raven (25), each of which produced a single, continuous sequence representing the bacterial chromosome, and most of which contained two medium-sized plasmids at a similar read depth to the chromosome. For each assembly, the length-filtered reads were mapped to the chromosome and plasmids Minimap2 (26) and mapped reads were retained for downstream analysis. The chromosomal and plasmid reads were then downsampled to 500× coverage using Filtlong (github.com/rrwick/Filtlong) and combined into a single high-quality assembly with Trycycler (27) incorporating 24 separate assemblies from six assemblers: Canu (28), Flye (22), Miniasm (24), NECAT (29), NextDenovo (30), and Raven (25). The final complete genome assembly was annotated with Bakta v1.9.1 (31). Full details of the individual steps for quality controls, kraken assignment and genome assembly are available in Supplementary File 1.

### Phylogeny of Coxiellaceae Family

The Genome Taxonomy Database Toolkit (GTDB-Tk) (32) classified this genome as Coxiellaceae. Representative isolates from the order Coxiellales (gtdb.ecogenomic.org/tree?r=o__Coxiellales) were used in comparative analyses to infer the relative location of Candidatus *P. cheracis* in the order. Given the large number of Coxiellales genomes in the GTDB database, the majority of which belong to the clinically important species *Coxiella burnetii*, the dataset was first dereplicated by Assembly Dereplicator (github.com/rrwick/Assembly-Dereplicator) to a Mash distance of 0.001, resulting in a representative set of 60 genomes capturing the available diversity of the Order. A phylogeny was inferred using GTDB-Tk (33), a marker gene-based tool for taxonomic classification and phylogenetic placement of prokaryotic genomes based on the Genome Taxonomy Database (GTDB) taxonomy system (34) was used to infer a phylogeny for the representative 60 genomes and to taxonomically classify our long-read assembly.

### Investigation of the complete genome for features of interest

The genome content of Candidatus *P. cheracis* was explored using publicly available tools. First, known antimicrobial resistance (AMR) determinants were screened with abritAMR (v1.0.14) with the AMRfinderPlus database (v2022-08-09.1) with no species flag and default parameters (35). Mob-typer, which is part of Mob-Suite, was used to identify plasmid replicons and mob genes with default parameters (36, 37). Phage defense mechanisms were identified with PADLOC v2 using the annotated gff file as input and default parameters (38).

Putative Dot/Icm effectors were screened in the complete genome with Bastion4 (39) and T4Sepp (40) tools. The predictions were compared, with hits for putative IS elements not included in further analysis due to the likelihood of being false positive results.

Components of the T4SS were identified with SECRET4 (41). T4SSs were also identified in selected, publicly available genomes that have been established to have functional T4SSs. These included two *C. burnetii* genomes as the type strains for the species (RSA493 accession: AE016828.3, and Dugway accession: CP000733.1), two genomes from the *Legionella* genus, *Legionella pneumophila* subsp*. pneumophila* str. Philadelphia 1 (accession: AE017354.1) and *Legionella longbeachae* NSW150 (accession:NC_013861.1)(42) and the Candidatus *Rickettsiella viridis* Ap-RA04 genome (accession: AP018005.1). We also screened representative isolates with complete genomes from *Coxiella* like-endosymbionts (C-LEs) (accessions: CP021379.1, CP011126.1, CP033868.1 and CP064834.1) with SECRET4 confirming the reported lack of a complete T4SS in these genomes (43).

Protein identity between the six genomes was undertaken with *blastp* using the Dot/Icm proteins from *C. burnetii* RSA493 (accession: AE016828.3) as a query sequence. For the three *Coxiella* genomes, the output was filtered to the top hit for each with a bitscore >100. A modified approach was undertaken for the two *Legionella* and Candidatus *Rickettsiella viridis* genomes due to the lower sequence homology. For these three genomes, the results were filtered to the loci that encode the T4SS and the hits with highest bitscore for each protein was included.

Visualization of the Candidatus *P. cheracis* genome with features of interest including phage defense, T4SS and putative Dot/Icm effectors was done in BRICK (github.com/esteinig/brick). The phylogenetic tree was visualized in R using ape (44) and ggtree (45). The R package gggenomes was used to plot the T4SS (github.com/thackl/gggenomes) and pheatmap was used to visualize the protein identity (github.com/raivokolde/pheatmap).

### Cultivation of bacterial and tissue culture cells

*C. burnetii* Nine Mile phase II (NMII) strain RSA439 clone 4 was axenically cultivated in liquid ACCM-2 or ACCM-2 agarose at 37°C in 5% CO_2_ and 2.5% O_2_ as previously described (46). When required chloramphenicol was added to *C. burnetii* cultures at 3 μg/ml. HeLa CCL-2 cells (ATCC, VA) were maintained in Dulbecco’s Modified Eagle’s Media (DMEM) supplemented with 10% heat inactivated fetal bovine serum at 37°C in 5% CO_2_.

### Construction of *C. burnetii* translocation reporter strains

The open reading frame encoding for the Candidatus *P. cheracis* CpeB homologue (*Pc*CpeB) was synthesized by IDT and cloned into pJB-CAT:BlaM (47) using BamHI and XhoI restriction enzymes. The resulting pBlaM-*Pc*CpeB was introduced into *C. burnetii* strains using a standard electroporation protocol (11). Following selection of transformants, expression of BlaM fusion proteins was confirmed using an anti-BlaM (1:1500, QED Biosciences). *C. burnetii* strains carrying either pJB-CAT:BlaM or pJB-CAT:BlaM:CBU0021 were used as negative and positive translocation controls (48).

### BlaM reporter translocation assay

Translocation assays were conducted as previously reported (17, 49). HeLa CCL2 cells were seeded in 96-well black, clear bottom tissue culture plates at a density of 10^4^ per well. Approximately 24 h later, cell monolayers were infected with *C. burnetii* strains at an MOI of 300. Following a 22 h incubation, wells were loaded with CCF2-AM substrate from the LiveBLAzer FRET loading kit (Thermo Fisher Scientific) and incubated at room temperature for 2 h. Wells were excited at 415 nm and the emission at 450 nm (blue) and 520 nm (green) was collected using a Clariostar fluorescence plate reader. Representative images were acquired using an Invitrogen EVOS™ FL Imaging System, images taken at 10× objective.

### Data availability

The complete genome and the long-read data are available at BioProject PRJNA1189271.

## RESULTS

### A putative new genus in the Coxiellaceae family

Here we describe the complete genome of a putative new genus within Coxiellaceae, Candidatus *C. cheraxi,* that is the likely causative agent of mass mortality events in *C. quadricarinatus*. The Candidatus *C. cheraxi* genome consisted of one chromosome (2,233,862 bp) and two smaller plasmids (59,783 and 34,021 bp). The chromosome is similar in size to previously sequenced *C. burnetii* isolates ranging from approximately 1.9 Mb to 2.2 Mb (50). To confirm that this organism is related to the original Candidatus *C. cheraxi* TO-98 organism, the 21 available sequence reads from TO-98 (6) were mapped to this genome with an average identify of 89%. This is a high level of similarity, even with the TO-98 dataset produced using older minion chemistry (R9.4.1), provided confidence that this genome is representative of the same species as TO-98.

The Candidatus *C. cheraxi* genome was classified as Coxiellaceae by GTDB-Tk (32), meaning that the novel genome could be classified to the family level but no further, suggesting that this genome represents a novel genus, supported by a Relative Evolutionary Divergence (RED) value of 0.79373 (32). Based on this data we propose to consider this Coxiellaceae as a novel genus: Candidatus *Paracoxiella cheracis*, also adopting the adjusted genus nomenclature (51).

A phylogenetic tree inferred from bacteria-specific marker genes placed Candidatus *P. cheracis* near genomes of other Coxiellaceae bacteria (**Figure 1**) isolated from similar aquatic ecological niches and distant to *C. burnetii* (largely associated with mammalian hosts) and the *Coxiella*-like endosymbionts (C-LEs) that are associated with ticks as hosts (**Figure 1**). Interestingly, Kraken2 classification of the chromosomal reads using a GTDB-based database identified multiple bacterial families in the read set. These include 10.6% Alteromonadaceae, 4.1% Vibrionaceae, 3.7% Enterobacteriaceae, 2.9% Burkholderiaceae, 2.5% Legionellaceae, 2.3% Moraxellaceae and only 8.5% Coxiellaceae (**Supplementary File 2**), which are all members of the Gammaproteobacteria class. This demonstrates that Kraken2 results for an isolate in a novel taxon not represented in the database can be misleading and may falsely suggest a mixed sample. However, assembly of the long-read data successfully recovered the bacterial chromosome and two plasmids.

**Figure 1.**
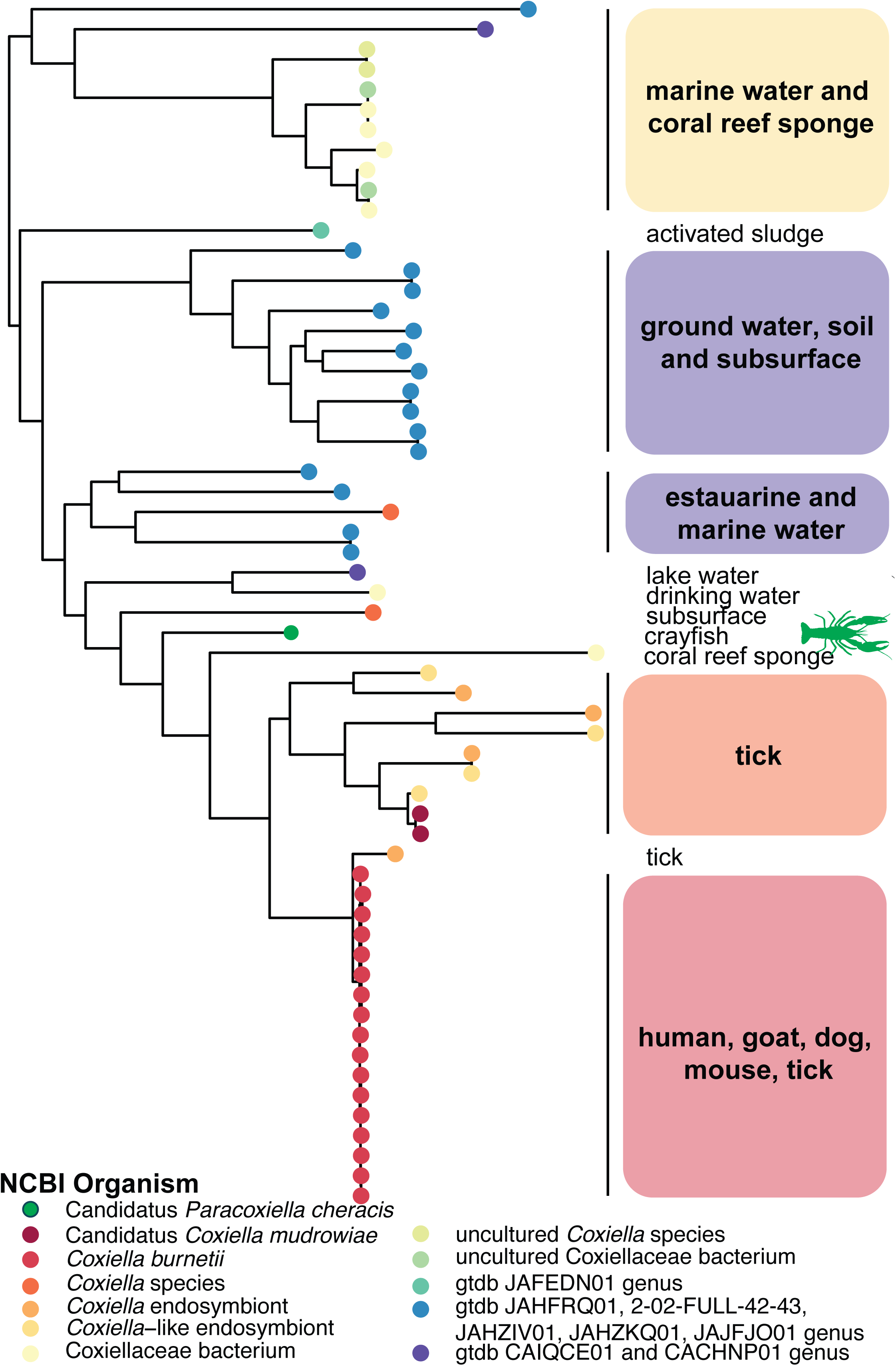
Phylogeny of Coxiellales Order demonstrating relationship to Candidatus *P. cheracis* genome. The inferred evolution of the Candidatus *P. cheracis* genome in representative isolates from Coxiellales Order (gtdb.ecogenomic.org/tree?r__oCoxiellales). The phylogeny was inferred using GTDB-Tk, a marker gene-based method of 120 domain-specific marker genes for bacteria (33). The tips of the tree are coloured by the NCBI organism for the Coxiellales lineages, and additional GTDB lineages at the genus level. The associated hosts or source from where the samples were collected are shown to the right of the tree. The Candidatus *P. cheracis* genome is indicated in green and by the crayfish icon (sourced from www.phylopic.org/).

The complete genome of Candidatus *P. cheracis* was explored for antimicrobial resistance, phage defense mechanisms, the Dot/Icm T4SS and putative effectors of this system (**Figure 2**). No known AMR mechanisms were identified, consistent with the AMR profiles of *C. burnetii* genomes. No replicon or mob genes were detected in the two plasmids. Further, these two plasmid sequences had no homology to publicly available data on NCBI. Several phage defense systems were identified on the chromosome in the novel genome. Two SoFic defense systems and the AbiE system, comprised of AbiEi and AbiEii components, were identified in the genome. The remaining ten defense systems identified were from the curated PADLOC database. Multiple putative effectors were identified (expanded upon in Predicted substrates of the *C. cheraxi* Dot/Icm secretion system), in addition to many copies and variants of a transposase that shares identity with a transposase from a symbiont of *Solemya velum*, the Atlantic awning clam [ref, accession: JRAA00000000] (52).

**Figure 2.**
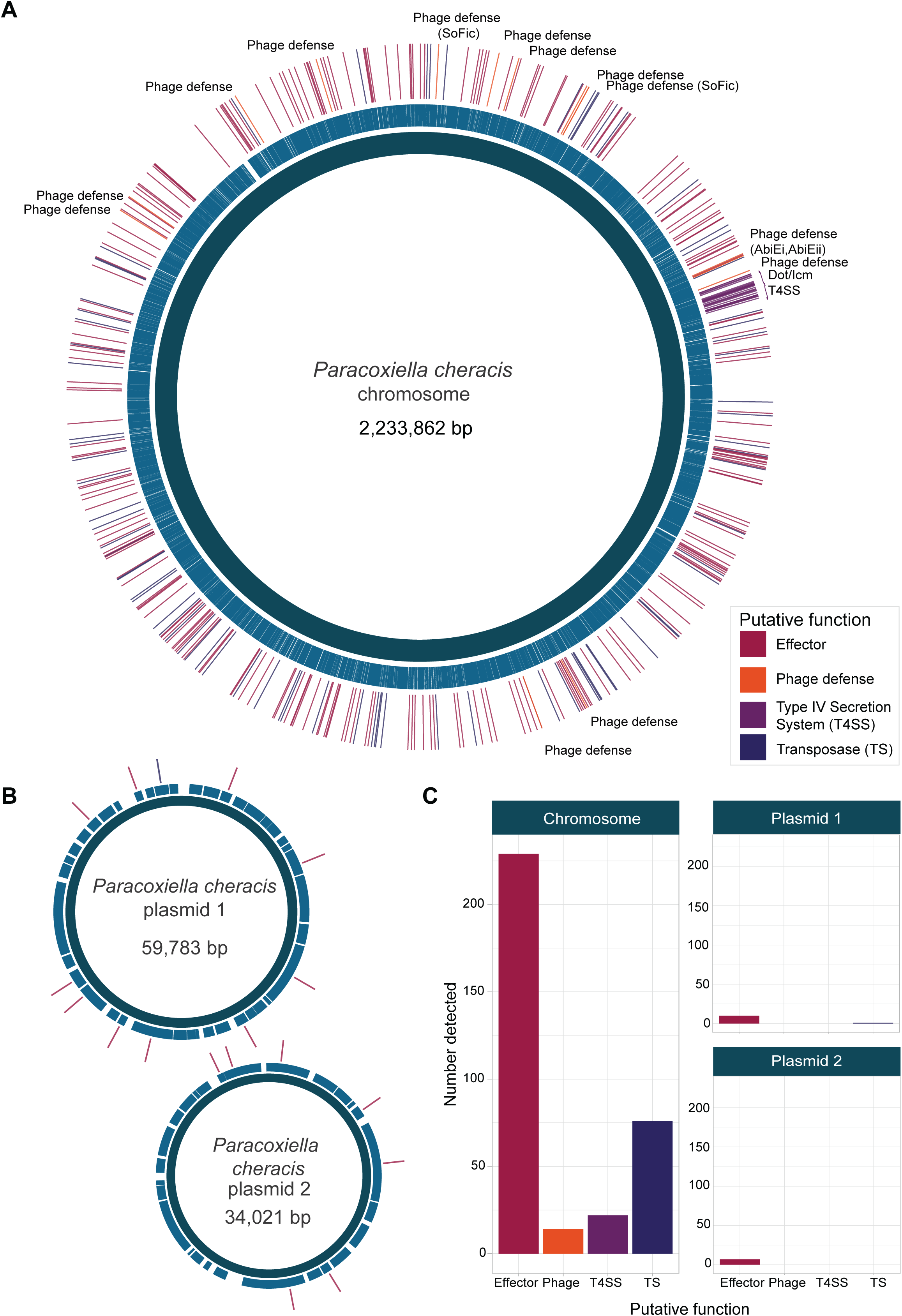
Characterization of the Candidatus *P. cheracis* genome. The complete genome showing A) the chromosome and B) two plasmids. The middle ring shows the coding sequence (CDS) on the chromosome and plasmids. Genes of interest are shown in the outer ring with colours indicating broad function (T4SS, multicopy transposase (TS), phage defense and putative effector). The total number of these four functional classes on each genetic element are shown on panel C.

### Detection and analysis of the Dot/Icm locus encoded by Candidatus *P. cheracis*

The Dot/Icm system was detected in the Candidatus *P. cheracis* genome, suggesting that the isolate has a functional T4SS. The site of integration into the chromosome and gene arrangement were similar to the two isolates of *C. burnetii* examined here (**Figure 3A**). The two *C. burnetii* genomes have additional genes between *icmT* and *icmQ,* some of which have been shown to be effector proteins (53), which are lacking in the Candidatus *C. cheraxi* genome. Additionally, *icmT* and *icmS* were encoded in the reverse orientation in the Candidatus *P. cheracis* genome compared to the two *C. burnetii* genomes. These two genes were in the same orientation as the two *Legionella* and Candidatus *R. viridis* genomes despite the Dot/Icm system being fragmented into multiple regions in these three genomes. Of note, the protein homology between 21 genes of the Dot/Icm system differed within the six genomes by species and genus (**Figure 3B**). The proteins from the *C. burnetii* Dugway isolate were homologous to the Dot/Icm proteins of *C. burnetii* RSA493 (>99% protein identity for all). In contrast, the protein identity for the Dot/Icm proteins in Candidatus *P. cheracis* varied between 52% (IcmG) and 86% (IcmB) identity relative to the *C. burnetii* RSA493 proteins. This was higher than the protein similarity of the Dot/Icm proteins from the two *Legionella* and the Candidatus *R. viridis* genomes which ranged between 24% (IcmX in *L. longbeachae*) to 67% (DotB for Candidatus *R. viridis*). As previously reported, the Dot/Icm system was not detected in the three C-LEs genomes.

**Figure 3.**
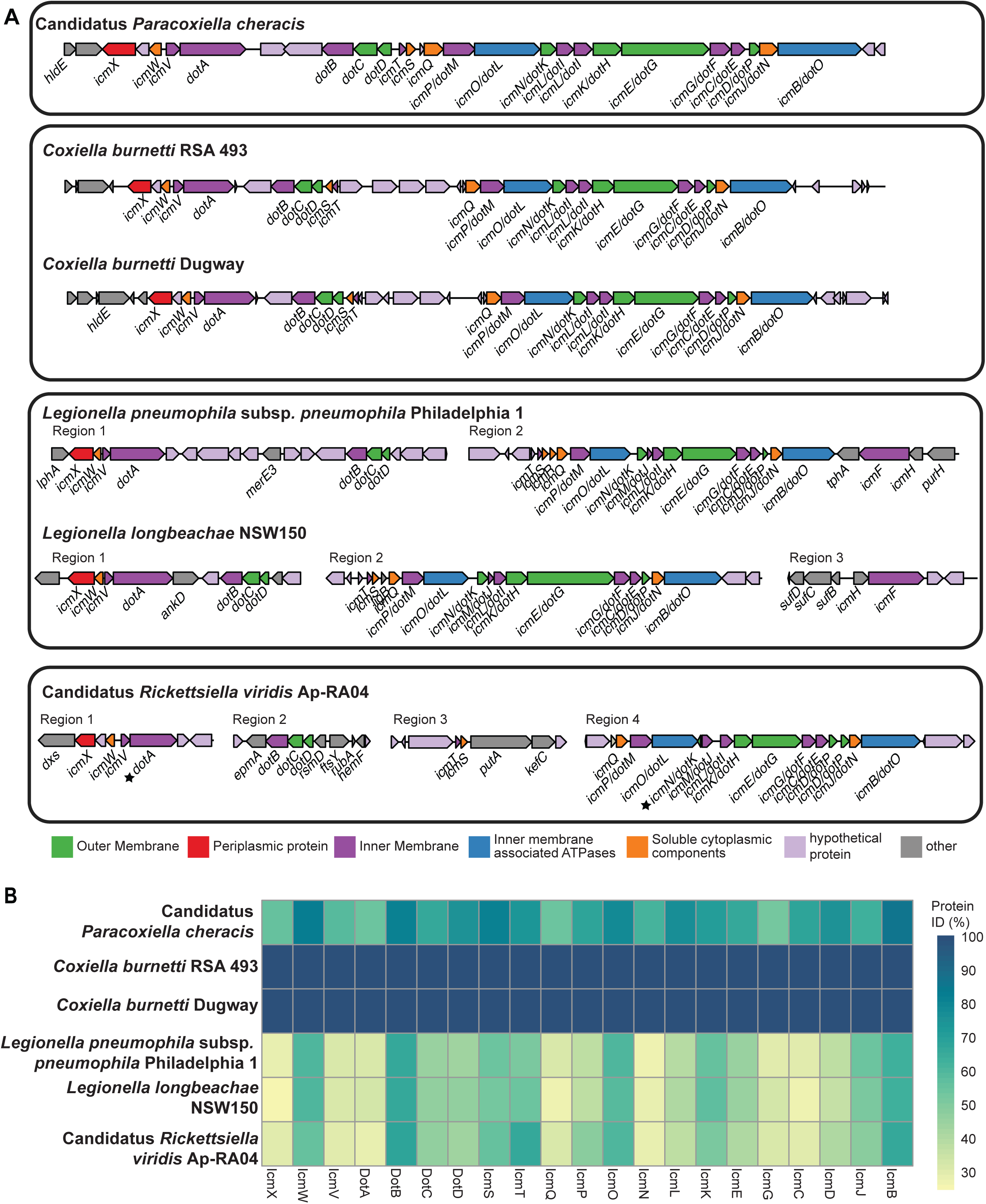
Structure and protein identity of the T4SS in the Candidatus *P. cheracis* genome. A) Characterization of the T4SS identified in Candidatus *P. cheracis, Coxiella burnetii*, *Legionella pneumophila, Legionella longbeachae* and Candidatus *Rickettsiella viridis.* Genes encoding the T4SS are colored by function within the system. B) Visualization of Dot/Icm protein identity relative to *Coxiella burnetii* RSA 493.

### Predicted substrates of the Candidatus *P. cheracis* Dot/Icm secretion system

Given the conservation of genes encoding for the Dot/Icm apparatus we hypothesized that Candidatus *P. cheracis* possesses a unique repertoire of putative T4SS substrates. To identify these proteins, two effector prediction pipelines, Bastion4 and T4SEpp, were employed. Bastion4, bacterial secretion effector predictor for T4SS, is an ensemble effector predictor based on six distinct machine learning models (39). More recently, T4SEpp was developed using an integrated pipeline that incorporates homology-based predictions with machine learning models to generate a prediction score for the likelihood of a protein being a T4SS substrate (40). Combined, these tools predicted 355 putative Dot/Icm substrates within the Candidatus *P. cheracis* genome. Bastion4 predicted 296 effectors and T4SEpp predicted 114 with 55 putative effectors predicted by both approaches (**Supplementary File 3**). Bastion4 predicted 117 instances of IS elements as putative effectors which are likely false positives bringing the pool of potential Dot/Icm effectors to 238. Of particular interest, 66 putative effectors (25 predicted by both tools) have no significant similarity to protein sequences found in NCBI.

### Candidatus *P. cheracis* CpeB, *Pc*CpeB, can be translocated by the *C. burnetii* Dot/Icm system

Each of the predicted Dot/Icm effectors were examined for similarity to known Dot/Icm effectors of *L. pneumophila* and *C. burnetii*. No putative effectors showed similarity within the *L. pneumophila* effector cohort however 12 putative effectors have similarity to putative and experimentally characterized *C. burnetii* effectors (**Table 1**).

**Table 1.**
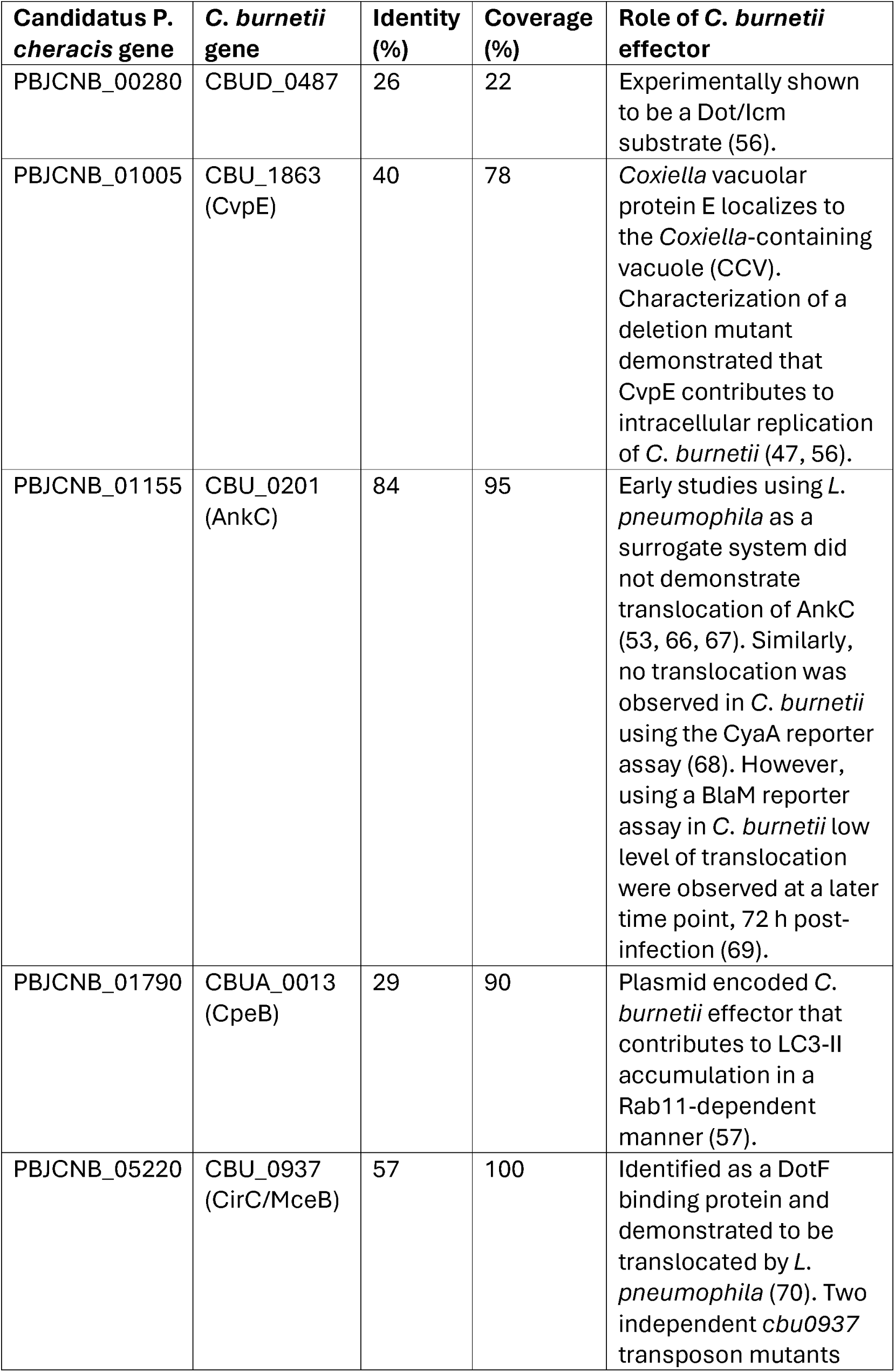
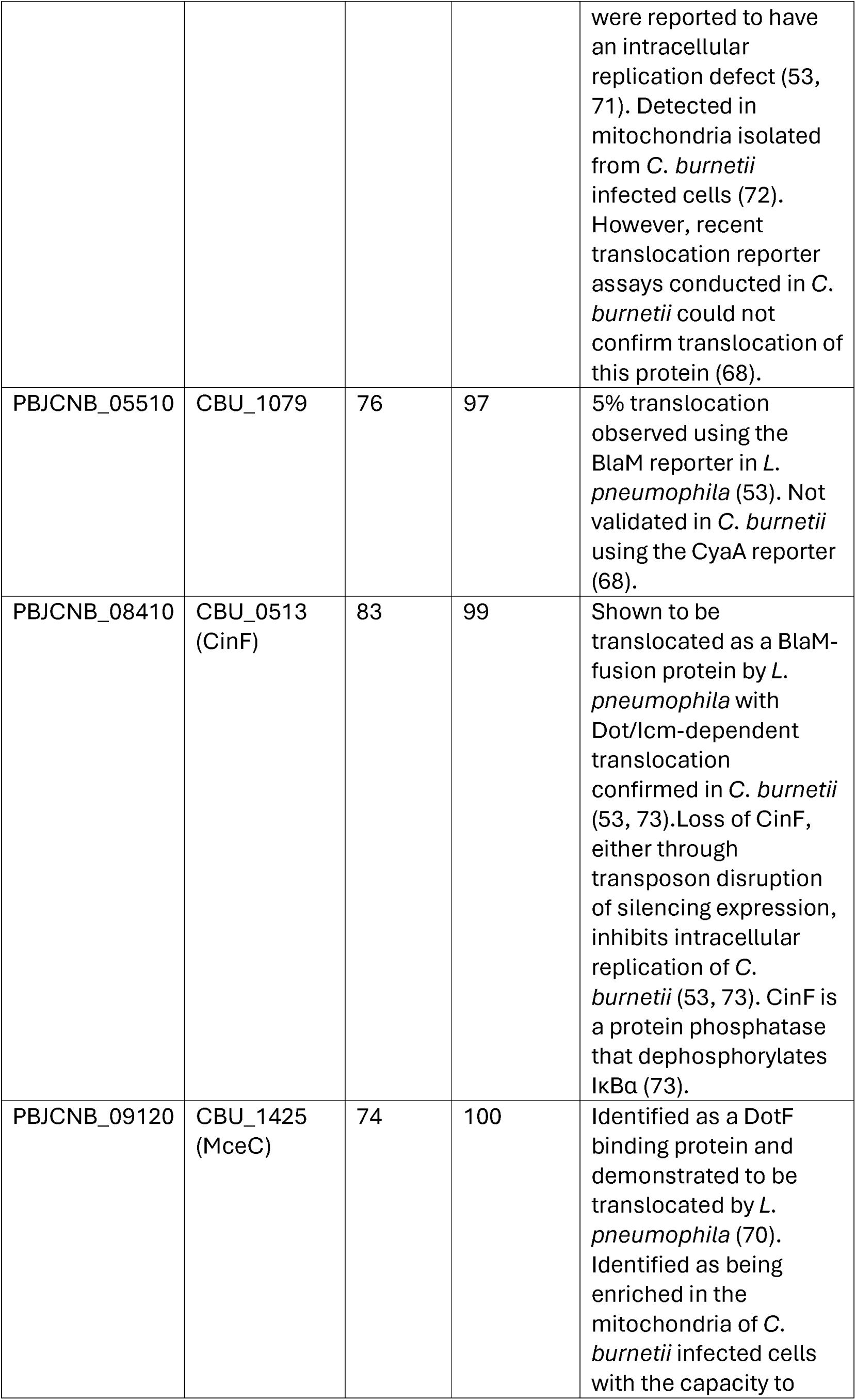
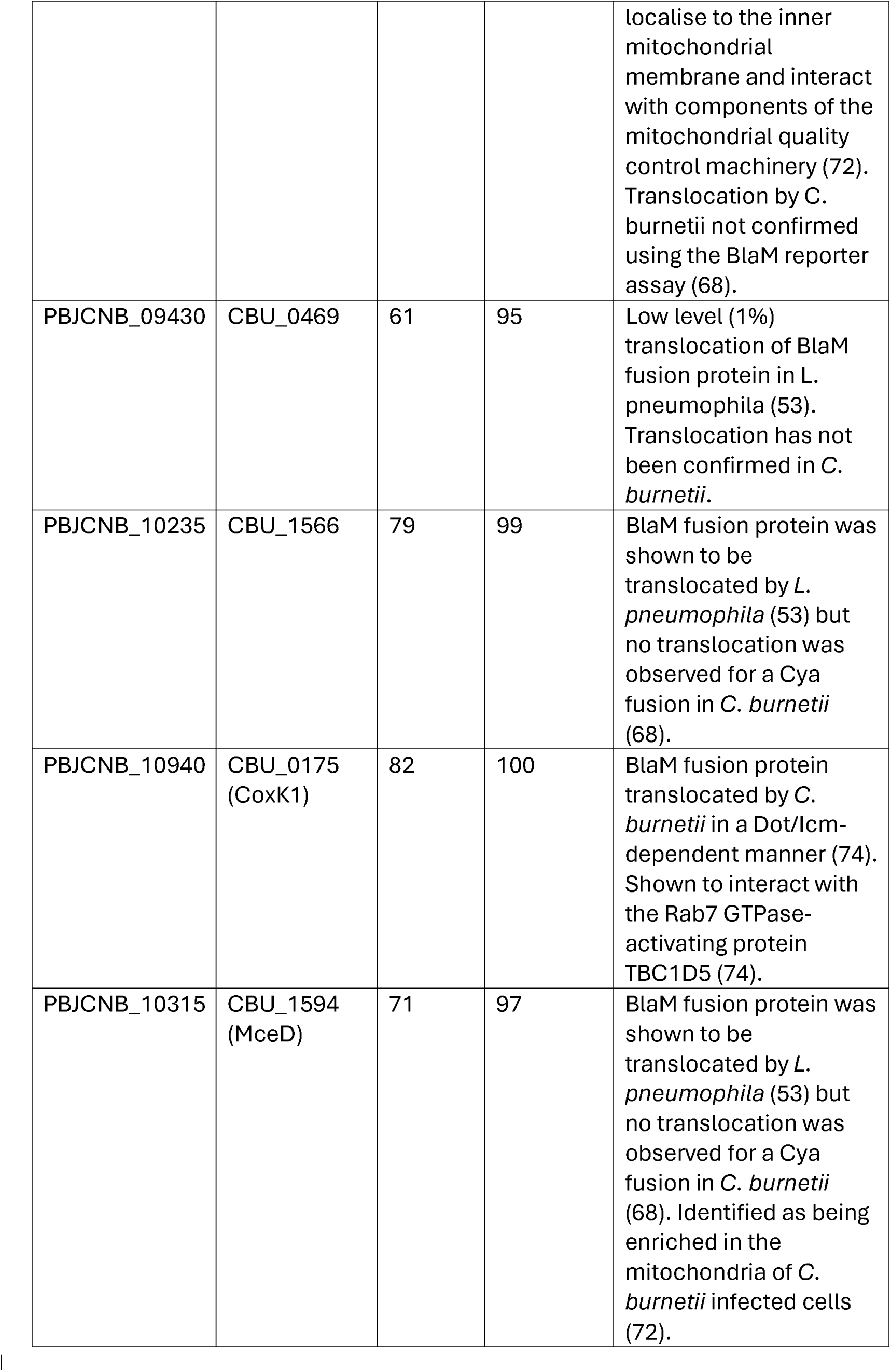
Candidatus P. cheracis proteins with similarity to C. burnetii Dot/Icm substrates.

| **Candidatus *P. cheracis* gene** | ***C. burnetii* gene** | **Identity (%)** | **Coverage (%)** | **Role of *C. burnetii* effector** |
| --- | --- | --- | --- | --- |
| PBJCNB_00280 | CBUD_0487 | 26 | 22 | Experimentally shown to be a Dot/Icm substrate (56). |
| PBJCNB_01005 | CBU_1863 (CvpE) | 40 | 78 | *Coxiella* vacuolar protein E localizes to the *Coxiella*-containing vacuole (CCV). Characterization of a deletion mutant demonstrated that CvpE contributes to intracellular replication of *C. burnetii* (47, 56). |
| PBJCNB_01155 | CBU_0201 (AnkC) | 84 | 95 | Early studies using *L. pneumophila* as a surrogate system did not demonstrate translocation of AnkC (53, 66, 67). Similarly, no translocation was observed in *C. burnetii* using the CyaA reporter assay (68). However, using a BlaM reporter assay in *C. burnetii* low level of translocation were observed at a later time point, 72 h post-infection (69). |
| PBJCNB_01790 | CBUA_0013 (CpeB) | 29 | 90 | Plasmid encoded *C. burnetii* effector that contributes to LC3-II accumulation in a Rab11-dependent manner (57). |
| PBJCNB_05220 | CBU_0937 (CirC/MceB) | 57 | 100 | Identified as a DotF binding protein and demonstrated to be translocated by *L. pneumophila* (70). Two independent *cbu0937* transposon mutants were reported to have an intracellular replication defect (53, 71). Detected in mitochondria isolated from *C. burnetii* infected cells (72). However, recent translocation reporter assays conducted in *C. burnetii* could not confirm translocation of this protein (68). |
| PBJCNB_05510 | CBU_1079 | 76 | 97 | 5% translocation observed using the BlaM reporter in *L. pneumophila* (53). Not validated in *C. burnetii* using the CyaA reporter (68). |
| PBJCNB_08410 | CBU_0513 (CinF) | 83 | 99 | Shown to be translocated as a BlaM-fusion protein by *L. pneumophila* with Dot/Icm-dependent translocation confirmed in *C. burnetii* (53, 73). Loss of CinF, either through transposon disruption or silencing expression, inhibits intracellular replication of *C. burnetii* (53, 73). CinF is a protein phosphatase that dephosphorylates IκBα (73). |
| PBJCNB_09120 | CBU_1425 (MceC) | 74 | 100 | Identified as a DotF binding protein and demonstrated to be translocated by *L. pneumophila* (70). Identified as being enriched in the mitochondria of *C. burnetii* infected cells with the capacity to localise to the inner mitochondrial membrane and interact with components of the mitochondrial quality control machinery (72). Translocation by *C. burnetii* not confirmed using the BlaM reporter assay (68). |
| PBJCNB_09430 | CBU_0469 | 61 | 95 | Low level (1%) translocation of BlaM fusion protein in *L. pneumophila* (53). Translocation has not been confirmed in *C. burnetii*. |
| PBJCNB_10235 | CBU_1566 | 79 | 99 | BlaM fusion protein was shown to be translocated by *L. pneumophila* (53) but no translocation was observed for a Cya fusion in *C. burnetii* (68). |
| PBJCNB_10940 | CBU_0175 (CoxK1) | 82 | 100 | BlaM fusion protein translocated by *C. burnetii* in a Dot/Icm-dependent manner (74). Shown to interact with the Rab7 GTPase-activating protein TBC1D5 (74). |
| PBJCNB_10315 | CBU_1594 (MceD) | 71 | 97 | BlaM fusion protein was shown to be translocated by *L. pneumophila* (53) but no translocation was observed for a Cya fusion in *C. burnetii* (68). Identified as being enriched in the mitochondria of *C. burnetii* infected cells (72). |

To test the feasibility of using *C. burnetii* as a surrogate bacterium to demonstrate Dot/Icm-dependent translocation of putative Candidatus *P. cheracis* effectors, the BlaM translocation reporter assay was employed. This assay is routinely used to demonstrate Dot/Icm substrate translocation into host cells and relies on bacterial delivery of a β-lactamase (BlaM) enzyme into the host cytosol where its activity can be measured by cleavage to the BlaM substrate CCF2-AM (11). Dot/Icm-dependent translocation of the Candidatus *P. cheracis* homologue of *C. burnetii* effector CpeB (*Pc*CpeB) was confirmed (**Figure 4A and B**). *C. burnetii* WT and Dot/Icm-deficient (*icmL::*Tn) strains were engineered to express BlaM alone (negative control), BlaM-CvpB, a well characterized *C. burnetii* Dot/Icm effector (positive control) (48, 54–56) or BlaM-*Pc*CpeB (**Fig 4C**). HeLa cells were infected with these *C. burnetii* strains for 22 h before addition of CCF2-AM and subsequent detection of fluorescence emission via both a fluorescence plate reader (**Fig 4A**) and microscopy (**Fig 4B**). BlaM-*Pc*CpeB was translocated in a Dot/Icm-dependent manner. Interestingly, comparison of the amino acid sequences of *C. burnetii* CpeB (*Cb*CpeB) and *Pc*CpeB showed only 29% identity however the region determined to mediate interaction between *Cb*CpeB and Rab11 shows greater conservation potentially indicating functional similarity (purple box, **Fig 4D**) (57).

**Figure 4.**
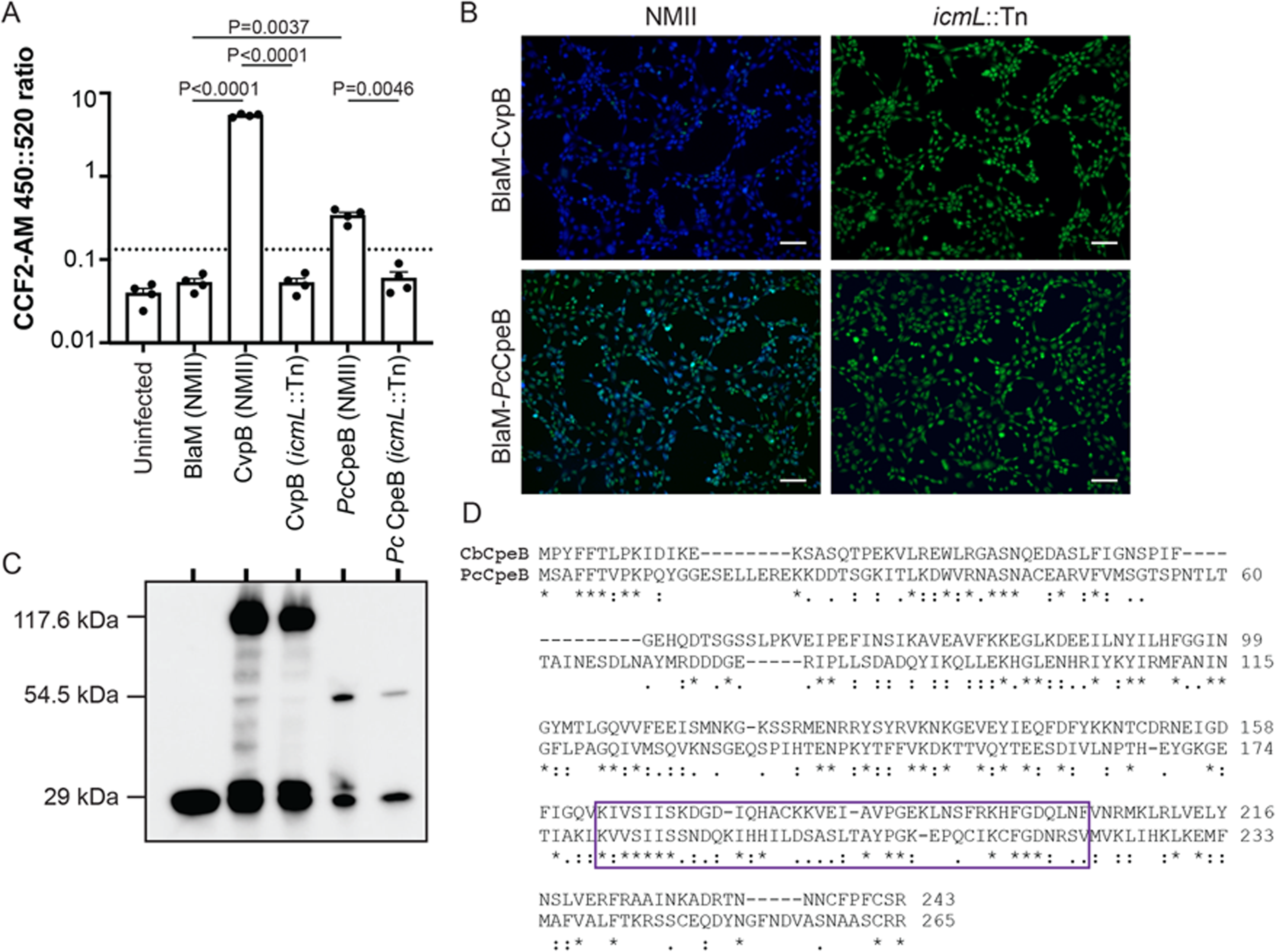
Candidatus *P. cheracis Pc*CpeB is a Dot/Icm substrate. HeLa cells were infected for 24 h with either wildtype (NMII) or Dot/Icm-deficient (*icmL*::Tn) *C. burnetii* expressing the indicated BlaM fusion proteins. **(A)** Change in the fluorescence emission ratio 450nm::520nm was measured as a readout of BlaM cleaving the CCF2-AM substrate (y-axis). The graph depicts this ratio from 4 independent experiments (circles) with the bar representing the mean and SEM ratio. The threshold for translocation positive was set to 2.5× the BlaM alone negative control (dotted line) and Šidák’s multiple comparisons test was used to determine P-values compared to the BlaM and between NMII and *icmL*::Tn strains expressing the same fusion proteins. **(B)** Representative fluorescent images of infected HeLa cells demonstrate the translocation of CvpB (*C. burnetii* effector positive control) and *Pc*CpeB by *C. burnetii* NMII but not *C. burnetii icmL*::Tn. Fluorescence intensity at 520 nm of uncleaved CCF2-AM is shown in green and cleaved CCF2-AM, at 450 nm is blue, scale bars represent 100 μm. **(C)** Immunoblot analysis of the indicated *C. burnetii* strains confirmed expression of BlaM (29 kDa), BlaM-CvpB (117.6 kDa) and BlaM-*Pc*CpeB (54.5 kDa). **(D)** ClustalOmega alignment showing the similarity between *C. burnetii* CpeB (*Cb*CpeB) and Candidatus *P. cheracis* CpeB (*Pc*CpeB). The two proteins share 26% identity (*) with additional conserved (:) and semi-conserved (.) substitutions. The purple boxed area represents amino acids 164 to 204 that were previously shown to mediate interaction of *Cb*CpeB with Rab11a (57).

## DISCUSSION

In the absence of live samples of Candidatus *P. cheracis* n.g. n. sp., progressing our understanding of this novel crayfish pathogen is limited to the exploration of genetic material. This report of the complete genome sequence of this pathogen provides both an opportunity to expand our understanding of the evolutionary relationships between members of the family Coxiellaceae and formulate hypotheses regarding the host-pathogen interactions mediated by this organism.

The phylogenetic placement, ANI to reference genomes, and Relative Evolutionary Divergence of Candidatus *P. cheracis* provides evidence that this novel genome represents a new genus within the Coxiellaceae family and is more distantly related to both *C. burnetii* and C-LEs than previously appreciated. Of note, the Dot/Icm T4SS detected in Candidatus *P. cheracis* was found to have inserted into the chromosome near the same genes as in *C. burnetii* genomes. From this, we hypothesize that the Dot/Icm T4SS may have been acquired in the most recent common ancestor of both Candidatus *P. cheracis, C. burnetii* and C-LE. Moreover, the loss of the Dot/Icm T4SS in the C-LEs which are associated with invertebrate hosts suggests that the Dot/Icm T4SS did not confer a fitness advantage to warrant being maintained in C-LEs, which are maternally inherited in ticks (19).

The Dot/Icm T4SS has co-evolved in these different genetic backgrounds, resulting in the observed differences in protein identity between Candidatus *P. cheracis* and the two *C. burnetii* reference genomes, with adaptation of the effectors to the different host niches. The observed differences in Dot/Icm protein identity between these genomes is greater than the average values recently reported for 58 *Legionella* species (58). However, the Dot/Icm components of *Legionella* with the greatest variation in protein identity were the same in the Candidatus *P. cheracis* (relative to *C. burnetii*), specifically IcmX, DotA and IcmG/DotF, consistent with exposure of these protein to hosts (58). In contrast, IcmB and DotB had the highest protein identity, which is consistent with previous work in *Legionella* species suggestive that these proteins are under functional selection (58).

Key challenges in the detection of obligate intracellular pathogens is the difficulty in culturing the organisms, and the lack of diagnostic targets that can detect these bacteria at low concentrations in often complex metagenomic samples. Here we identified that Candidatus *P. cheracis* possesses a high copy transposase that had homology to a symbiont of *Solemya velum*, the Atlantic awning clam. Future efforts to screen for the presence of Candidatus *P. cheracis* could be facilitated by PCR amplification of the transposase in aquatic samples. This would be akin to diagnostic efforts of members of the *Bordetella* genus (59). *Bordetella pertussis,* the causative agent of whooping cough, is commonly identified through the PCR of high copy number insertion sequences (IS) with the primary target IS481. This would provide a potentially non-invasive, sensitive screening strategy for ongoing surveillance of the crayfish farms that would facilitate the prevention and absence of this pathogen in aquaculture environments. Additionally, if proven as a sensitive strategy for detection of Candidatus *P. cheracis*, this PCR approach could also be employed to test different aquatic environments for the presence of this Coxiellaceae genus.

Comparative genomics of 65 *Legionella* species has reported an expansive repertoire of putative Dot/Icm effector proteins (60). While many species possess putative effectors with specific eukaryotic domains, such as GTPase, F-box and SET domains, these motifs are encoded within different proteins that are not orthologous (60). Interestingly, only 8 Dot/Icm substrates are conserved among these 65 species of *Legionella* (60). Using two distinct *in silico* tools, we have developed a list of 238 putative Dot/Icm effectors of Candidatus *P. cheracis*. Unlike *Legionella* species, very few putative Dot/Icm effectors encoded by Candidatus *P. cheracis* possess similarity to eukaryotic domains and a large proportion, ∼28%, are novel. We observed 12 predicted effectors with similarity to known or putative *C. burnetii* Dot/Icm effectors, including CpeB and CvpE which have been shown to make important contributions to the intracellular success of *C. burnetii* (48, 54, 56, 57). Further expanding the collection of sequenced Coxiellaceae genomes will facilitate greater comparison of the effector repertoire in this family, allowing us to determine whether some of these 12 common effectors act as “core” Coxiellaceae effectors that facilitate their characteristic interaction with eukaryotic cells. We were able to demonstrate Dot/Icm-dependent translocation of Candidatus *P. cheracis* CpeB (*Pc*CpeB) using *C. burnetii* as a surrogate host. In the absence of methodology for the isolation and cultivation of Candidatus *P. cheracis,* the success of this approach provides an alternative strategy for developing our understanding of the host-pathogen interactions mediated by Candidatus *P. cheracis*.

The full genome sequence of Candidatus *P. cheracis* has highlighted potential screening or diagnostic targets and unveiled likely pathogenesis strategies. In addition, the genome sequence shows that Candidatus *P. cheracis* encodes for Mip, a peptidyl-prolyl cis-trans isomerase that has been shown to be essential for intracellular replication and pathogenesis of *C. burnetii* (61). Candidatus *P. cheracis* Mip shares 66% identity with *C. burnetii* Mip and these proteins are conserved within the PPIase catalytic domain which suggests that Mip inhibitors developed by Debowski et al., may inhibit Candidatus *P. cheracis* Mip (61). Treatment with rationally designed small molecule inhibitors of Mip may represent a plausible intervention measure for future outbreaks of Candidatus *P. cheracis* in aquaculture facilities.

Here, we have further demonstrated the effective use of ONT sequencing to reconstruct the genome of a novel intracellular pathogen - a member of a family that is difficult to culture in standard laboratory conditions - directly from its host. The complete genome of a chromosome and two plasmids enabled the exploration of a previously unexplored pathogen that is important for Australian aquaculture. ONT was advantageous here for the capture of the two plasmids, elements that are often missed by short-read metagenomic binning approaches but can be critical for virulence (62, 63) and for dealing with the significant number of IS elements throughout this genome (64). The detection of a T4SS and cohort of novel effector proteins that likely facilitate intracellular success of the pathogen provides the first insights into the host pathogen interactions of this novel pathogen. ONT sequencing has shown promise in recent years for outbreak investigations due to its low-cost, portability, minimal sample preparation, and rapid, real-time data output (65). If such an outbreak was to occur today, in addition to traditional histopathology, ONT metagenomics would enhance efforts to detect and characterize unknown emerging pathogens.

## ACKNOWLEDGEMENTS

DJI is supported by a National Health and Medical Research Council (NHMRC) Investigator Grant (GNT1195210). CJW and RRW are funded through the NHMRC (GNT1194325). GRS is the recipient of an Australian Government Research Training Program Scholarship and Monash Graduate Excellence Scholarship. Research in HJNs laboratory was funded by the NHMRC (GNT2010841). The funders had no role in study design, data collection and analysis, decision to publish or preparation of the manuscript.

## SUPPLEMENTARY FILES

**Supplementary File 1**. Workflow for quality controls, kraken assignment and genome assembly.

**Supplementary File 2.** The Kraken2 classification of the chromosomal reads using a GTDB-based database.

**Supplementary File 3.** Predicted Dot/Icm effector proteins based on Bastion4 and T4Sepp tools.

## Notes

### Competing Interest Statement

The authors have declared no competing interest.

### Summary of Updates

This revision includes the three supplementary files.

