## Supplementary File 1 for "The complete genome sequence of the crayfish pathogen Candidatus *Paracoxiella cheracis* n.g. n.sp. provides insight into pathogenesis and the phylogeny of the Coxiellaceae family"

# Sequencing data

Two runs:
* DAMGPR0146_01 (MinION), where this sample (DMG2212273_2) was one of two barcodes.
  * Sequenced earlier (Jan 2023) and so used the older 4 kHz sampling rate.
  * Used adaptive sampling based on a _Coxiella_ reference genome.
* ONT_231215B (PromethION), where this was the only sample on the run.
  * Sequenced later (Dec 2023) and so used the newer 5 kHz sampling rate.
  * No adaptive sampling.


# ONT basecalling

## DMG2212273_2

For DMG2212273_2, I started with the fast5s that were already demultiplexed during the run, converted them to pod5, then basecalled. I used the `dna_r10.4.1_e8.2_400bps_sup@v4.1.0` model, which is the last one (I think) for 4kHz data. Done on OnION:
```bash
model=dna_r10.4.1_e8.2_400bps_sup@v4.1.0
kit=SQK-NBD114-96
run=DMG2212273_2

cd /data/"$run"
pod5 convert fast5 fast5/*.fast5 --output pods5/ --one-to-one ./fast5s/

mkdir "$model"
~/Tools/dorado-0.5.0-linux-x64/bin/dorado basecaller --recursive ~/Tools/dorado-0.5.0-linux-x64/models/"$model" pod5s --kit-name "$kit" > "$model"/reads.bam
cd "$model"
~/Tools/dorado-0.5.0-linux-x64/bin/dorado summary reads.bam > sequencing_summary.txt
```

I then combined all reads that were >1 kbp (no other QC because these were pass reads):
```bash
cd /home/wickr/operation_crayfish/DMG2212273_2/dna_r10.4.1_e8.2_400bps_sup@v4.1.0
samtools fastq -T '*' reads.bam | tr '\t' ' ' | paste - - - - > temp

# SPLIT BY CHANNEL
# adaptive sampling (bad): 1-412
# no adaptive sampling (good): 413-512

cat temp | grep -P " ch:i:\d " | tr '\t' '\n' > as.fastq
cat temp | grep -P " ch:i:\d\d " | tr '\t' '\n' >> as.fastq
cat temp | grep -P " ch:i:[123]\d\d " | tr '\t' '\n' >> as.fastq
cat temp | grep -P " ch:i:40\d " | tr '\t' '\n' >> as.fastq
cat temp | grep -P " ch:i:41[012] " | tr '\t' '\n' >> as.fastq

cat temp | grep -P " ch:i:41[3456789] " | tr '\t' '\n' > no_as.fastq
cat temp | grep -P " ch:i:4[23456789]\d " | tr '\t' '\n' >> no_as.fastq
cat temp | grep -P " ch:i:5\d\d " | tr '\t' '\n' >> no_as.fastq
rm temp

grep -oP " ch:i:\d+ " as.fastq | sort | uniq
grep -oP " ch:i:\d+ " no_as.fastq | sort | uniq

filtlong --min_length 1000 as.fastq | paste - - - - | sort | tr '\t' '\n' | pigz -p8 > ../../DMG2212273_2_as.fastq.gz
filtlong --min_length 1000 no_as.fastq | paste - - - - | sort | tr '\t' '\n' | pigz -p8 > ../../DMG2212273_2_no_as.fastq.gz
pigz -p8 *.fastq
```

## ONT_231215B

For ONT_231215B, I basecalled all pod5s from the entire run. I used the `dna_r10.4.1_e8.2_400bps_sup@v4.3.0` model, which is the last one for 5kHz data. Done on OnION:
```bash
model=dna_r10.4.1_e8.2_400bps_sup@v4.3.0
kit=SQK-RBK114-96
run=ONT_231215B

cd /data/"$run"
mkdir "$model"
~/Tools/dorado-0.5.0-linux-x64/bin/dorado basecaller --recursive ~/Tools/dorado-0.5.0-linux-x64/models/"$model" no_sample --kit-name "$kit" > "$model"/reads.bam
cd "$model"
~/Tools/dorado-0.5.0-linux-x64/bin/dorado summary reads.bam > sequencing_summary.txt
```

I then combined all reads that were >1 kbp and took the best 95%:
```bash
cd /home/wickr/operation_crayfish/ONT_231215B/dna_r10.4.1_e8.2_400bps_sup@v4.3.0
samtools fastq -T '*' reads.bam | tr '\t' ' ' > temp1.fastq
filtlong --min_length 1000 temp1.fastq >> temp2.fastq
filtlong --keep_percent 95 temp2.fastq | paste - - - - | sort | tr '\t' '\n' | pigz -p8 > ../../ONT_231215B.fastq.gz
rm temp1.fastq temp2.fastq
```

```bash
cd ~/operation_crayfish/ONT_231215B
mkdir reads
mv ONT* DMG* reads
```


# Kraken2

```bash
conda activate kraken
kraken2 --db ~/gtdb_r214_256gb --report reads/DMG2212273_2_as.report --threads 32 reads/DMG2212273_2_as.fastq.gz > reads/DMG2212273_2_as.kraken
kraken2 --db ~/gtdb_r214_256gb --report reads/DMG2212273_2_no_as.report --threads 32 reads/DMG2212273_2_no_as.fastq.gz > reads/DMG2212273_2_no_as.kraken
kraken2 --db ~/gtdb_r214_256gb --report reads/ONT_231215B.report --threads 32 reads/ONT_231215B.fastq.gz > reads/ONT_231215B.kraken
```


# Combine reads

For assembly, I'd like to just use the no-adaptive-sampling reads:
```bash
cd /home/wickr/operation_crayfish
zcat reads/DMG2212273_2_no_as.fastq.gz reads/ONT_231215B.fastq.gz | paste - - - - | sort | tr '\t' '\n' | pigz -p8 > reads/combined.fastq.gz
```

Since the PromethION run had much more yield, then combined reads were about 96% PromethION (5 kHz, sup@v4.3.0) and 4% MinION (4 kHz, sup@v4.1.0). So for later steps that care (Medaka), I can assume sup@v4.3.0.


# Initial assemblies

To see what the data looks like, I'm trying a few different assemblers.


## Flye

```bash
conda activate flye
flye --meta --nano-hq reads/combined.fastq.gz -o flye -t 64
any2fasta flye/assembly_graph.gfa > flye/assembly_graph.fasta
```


## Canu

To speed things up, I tried fiddling with some parameters:
* `rawErrorRate=0.2 correctedErrorRate=0.1` to account for modern ONT accuracy
* `minReadLength=4000 minOverlapLength=1000` because I have a lot of depth on the genome of interest
* `maxInputCoverage=10000 corOutCoverage=10000 corMhapSensitivity=high corMinCoverage=0 redMemory=32 oeaMemory=32 batMemory=200` are the Canu-doc-recommended metagenome parameters

```bash
canu -p canu -d canu -fast genomeSize=20000000 useGrid=false rawErrorRate=0.2 correctedErrorRate=0.1 minReadLength=4000 minOverlapLength=1000 minThreads=16 maxThreads=64 maxInputCoverage=10000 corOutCoverage=10000 corMhapSensitivity=high corMinCoverage=0 redMemory=32 oeaMemory=32 batMemory=200 -nanopore reads/combined.fastq.gz > canu.out 2>&1
```

```bash
~/Programs/Trycycler/scripts/canu_trim.py canu.fasta > canu_trimmed.fasta
mv canu_trimmed.fasta canu.fasta
```


## miniasm

```bash
cd /home/wickr/operation_crayfish
mkdir miniasm
miniasm_and_minipolish.sh reads/combined.fastq.gz 64 > miniasm/miniasm.gfa 2> miniasm/miniasm.out
any2fasta miniasm/miniasm.gfa > miniasm/miniasm.fasta
```

## Raven

```bash
cd /home/wickr/operation_crayfish
mkdir raven
raven --threads 64 --disable-checkpoints --graphical-fragment-assembly raven/raven.gfa reads/combined.fastq.gz > raven/raven.fasta 2> raven/raven.out
# I manually made raven_simple.gfa with just what I think are bacterial components.
any2fasta raven/raven_simple.gfa > raven/raven_simple.fasta
```


## metaMDBG

```bash
conda activate metamdbg
metaMDBG asm metamdbg reads/combined.fastq.gz -t 64 > metamdbg.out 2>&1
```


# Extract chromosome reads

For Flye, metaMDBG, miniasm and Raven, I got a nice chromosome for the Coxiellaceae genome. I'll use these as references to pull out the reads for this chromosome so I can do a Trycycler assembly.

My awk filter (`$3 < 10 && $2-$4 < 10`) required the entirety of the read to be aligned, allowing for a few unaligned bases at the start/end.

```bash
cd /home/wickr/operation_crayfish/Coxiellaceae_chromosome
r=/home/wickr/operation_crayfish/reads/combined.fastq.gz

minimap2 -c -t 64 flye.fasta "$r" | awk '$3 < 10 && $2-$4 < 10' | cut -f1 > flye.reads
minimap2 -c -t 64 metamdbg.fasta "$r" | awk '$3 < 10 && $2-$4 < 10' | cut -f1 > metamdbg.reads
minimap2 -c -t 64 miniasm.fasta "$r" | awk '$3 < 10 && $2-$4 < 10' | cut -f1 > miniasm.reads
minimap2 -c -t 64 raven.fasta "$r" | awk '$3 < 10 && $2-$4 < 10' | cut -f1 > raven.reads
cat flye.reads metamdbg.reads miniasm.reads raven.reads | sort | uniq > chromosome.reads
```

Of the 2558669 reads in `combined.fastq.gz`, about 44% matched the chromosome sequence.

I made this `sort_reads.py` script and put it in `~/operation_crayfish/scripts`:
```python
#!/usr/bin/env python3

import gzip
import sys

input_filename = sys.argv[1]
names_filename = sys.argv[2]
match_out_filename = sys.argv[3]
no_match_out_filename = sys.argv[4]

read_names = set()
with open(names_filename, 'rt') as f:
    for line in f:
        read_names.add(line.strip())

with gzip.open(input_filename, 'rt') as f, open(match_out_filename, 'wt') as match_out, open(no_match_out_filename, 'wt') as no_match_out:
    for header in f:
        seq = next(f)
        _ = next(f)
        qual = next(f)
        name = header[1:].split(' ')[0]
        out_f = match_out if name in read_names else no_match_out
        out_f.write(f'{header}{seq}+\n{qual}')
```

Extract Coxiellaceae chromosome reads into their own FASTQ:
```bash
cd ~/operation_crayfish/reads
../scripts/sort_reads.py combined.fastq.gz ../Coxiellaceae_chromosome/chromosome.reads chromosome.fastq other.fastq
```

This produced over 4 Gbp of chromosomal reads, so I Filtlonged the set down to 500x depth (1.1 Gbp):
```bash
filtlong --target_bases 1116928500 chromosome.fastq > chromosome_filtlong.fastq
pigz -p16 *.fastq
```

This resulted in 103244 reads with an N50 >10 kbp.


# Trycycler for Coxiellaceae chromosome

```bash
cd /home/wickr/operation_crayfish/chromosome
trycycler subsample --reads chromosome_filtlong.fastq.gz --out_dir read_subsets --count 24 --genome_size 2233857 2> subsample.out
```

Assembly following the [extra-thorough instructions](https://github.com/rrwick/Trycycler/wiki/Generating-assemblies#extra-thorough-assembly):
```bash
cd /home/wickr/operation_crayfish/chromosome
mkdir assemblies
genome_size=2233857

for i in 01 07 13 19; do
    ~/programs/canu/build/bin/canu -p canu -d canu_temp -fast genomeSize="$genome_size" useGrid=false maxThreads=32 maxMemory=120 -nanopore read_subsets/sample_"$i".fastq
    canu_trim.py canu_temp/canu.contigs.fasta > assemblies/assembly_"$i".fasta
    rm -rf canu_temp
done
for i in 02 08 14 20; do
    flye --nano-hq read_subsets/sample_"$i".fastq --threads 32 --out-dir flye_temp
    cp flye_temp/assembly.fasta assemblies/assembly_"$i".fasta
    cp flye_temp/assembly_graph.gfa assemblies/assembly_"$i".gfa
    rm -r flye_temp
done
for i in 03 09 15 21; do
    miniasm_and_minipolish.sh read_subsets/sample_"$i".fastq 32 > assemblies/assembly_"$i".gfa
    any2fasta assemblies/assembly_"$i".gfa > assemblies/assembly_"$i".fasta
done
for i in 04 10 16 22; do
    ~/programs/NECAT/Linux-amd64/bin/necat.pl config config.txt
    realpath read_subsets/sample_"$i".fastq > read_list.txt
    sed -i "s/PROJECT=/PROJECT=necat/" config.txt
    sed -i "s/ONT_READ_LIST=/ONT_READ_LIST=read_list.txt/" config.txt
    sed -i "s/GENOME_SIZE=/GENOME_SIZE="$genome_size"/" config.txt
    sed -i "s/THREADS=4/THREADS=32/" config.txt
    ~/programs/NECAT/Linux-amd64/bin/necat.pl bridge config.txt
    cp necat/6-bridge_contigs/polished_contigs.fasta assemblies/assembly_"$i".fasta
    rm -r necat config.txt read_list.txt
done
for i in 05 11 17 23; do
    echo read_subsets/sample_"$i".fastq > input.fofn
    cp ~/programs/NextDenovo/doc/run.cfg nextdenovo_run.cfg
    sed -i "s/genome_size = 1g/genome_size = "$genome_size"/" nextdenovo_run.cfg
    sed -i "s/parallel_jobs = 20/parallel_jobs = 1/" nextdenovo_run.cfg
    sed -i "s/read_type = clr/read_type = ont/" nextdenovo_run.cfg
    sed -i "s/pa_correction = 3/pa_correction = 1/" nextdenovo_run.cfg
    sed -i "s/correction_options = -p 15/correction_options = -p 32/" nextdenovo_run.cfg
    sed -i "s/-t 8/-t 32/" nextdenovo_run.cfg
    nextDenovo nextdenovo_run.cfg
    cp 01_rundir/03.ctg_graph/nd.asm.fasta nextdenovo_temp.fasta
    rm -r 01_rundir nextdenovo_run.cfg input.fofn
    echo read_subsets/sample_"$i".fastq > lgs.fofn
    cat ~/programs/NextPolish/doc/run.cfg | grep -v "sgs" | grep -v "hifi" > nextpolish_run.cfg
    sed -i "s/parallel_jobs = 6/parallel_jobs = 1/" nextpolish_run.cfg
    sed -i "s/multithread_jobs = 5/multithread_jobs = 32/" nextpolish_run.cfg
    sed -i "s|genome = ./raw.genome.fasta|genome = nextdenovo_temp.fasta|" nextpolish_run.cfg
    sed -i "s|-x map-ont|-x map-ont -t 32|" nextpolish_run.cfg
    nextPolish nextpolish_run.cfg
    cp 01_rundir/genome.nextpolish.fasta assemblies/assembly_"$i".fasta
    rm -r 01_rundir pid*.log.info nextpolish_run.cfg lgs.fofn nextdenovo_temp.fasta
done
for i in 06 12 18 24; do
    raven --threads 32 --disable-checkpoints --graphical-fragment-assembly assemblies/assembly_"$i".gfa read_subsets/sample_"$i".fastq > assemblies/assembly_"$i".fasta
done
```

```bash
cd /home/wickr/operation_crayfish/chromosome
trycycler cluster --assemblies assemblies/*.fasta --reads chromosome_filtlong.fastq.gz --out_dir trycycler --threads 48
trycycler reconcile --reads chromosome_filtlong.fastq.gz --cluster_dir trycycler/cluster_001
trycycler msa --threads 32 --cluster_dir trycycler/cluster_001
gunzip -c chromosome_filtlong.fastq.gz > trycycler/cluster_001/4_reads.fastq  # in place of trycycler partition
trycycler consensus --cluster_dir trycycler/cluster_001
```

```bash
cd /home/wickr/operation_crayfish/chromosome
for c in trycycler/cluster_*; do
    medaka_consensus -i "$c"/4_reads.fastq -d "$c"/7_final_consensus.fasta -o "$c"/medaka -m r1041_e82_400bps_sup_v4.3.0 -t 12
    mv "$c"/medaka/consensus.fasta "$c"/8_medaka.fasta
    rm -r "$c"/medaka "$c"/*.fai "$c"/*.mmi  # clean up
done
```


# Extract plasmid reads

There are two medium-sized plasmids which appeared in the Flye and miniasm assemblies (and one in the Raven assembly).

My awk filter (`$3 < 10 && $2-$4 < 10`) required the entirety of the read to be aligned, allowing for a few unaligned bases at the start/end.

```bash
cd /home/wickr/operation_crayfish/Coxiellaceae_plasmids
r=/home/wickr/operation_crayfish/reads/other.fastq.gz

minimap2 -c -t 64 flye_1.fasta "$r" | awk '$3 < 10 && $2-$4 < 10' | cut -f1 > flye_1.reads
minimap2 -c -t 64 miniasm_1.fasta "$r" | awk '$3 < 10 && $2-$4 < 10' | cut -f1 > miniasm_1.reads
minimap2 -c -t 64 flye_2.fasta "$r" | awk '$3 < 10 && $2-$4 < 10' | cut -f1 > flye_2.reads
minimap2 -c -t 64 miniasm_2.fasta "$r" | awk '$3 < 10 && $2-$4 < 10' | cut -f1 > miniasm_2.reads
minimap2 -c -t 64 raven_2.fasta "$r" | awk '$3 < 10 && $2-$4 < 10' | cut -f1 > raven_2.reads
cat flye_1.reads miniasm_1.reads flye_2.reads miniasm_2.reads raven_2.reads | sort | uniq > plasmid.reads
```

Extract Coxiellaceae chromosome reads into their own FASTQ:
```bash
cd ~/operation_crayfish/reads
mv other.fastq.gz temp.fastq.gz
../scripts/sort_reads.py temp.fastq.gz ../Coxiellaceae_plasmids/plasmid.reads plasmid.fastq other.fastq
rm temp.fastq.gz
```

This produced over 128 Mbp of plasmid reads, so I Filtlonged the set down to 500x depth (1.1 Gbp):
```bash
filtlong --target_bases 46901500 plasmid.fastq > plasmid_filtlong.fastq
pigz -p16 *.fastq
```

This resulted in 5296 reads with an N50 >8 kbp.


# Trycycler for plasmids

```bash
cd /home/wickr/operation_crayfish/plasmids
trycycler subsample --reads plasmid_filtlong.fastq.gz --out_dir read_subsets --count 24 --genome_size 93803
```

Assembly following the [extra-thorough instructions](https://github.com/rrwick/Trycycler/wiki/Generating-assemblies#extra-thorough-assembly):
```bash
cd /home/wickr/operation_crayfish/plasmids
mkdir assemblies
genome_size=93803

for i in 01 07 13 19; do
    ~/programs/canu/build/bin/canu -p canu -d canu_temp -fast genomeSize="$genome_size" useGrid=false minThreads=32 maxThreads=32 maxMemory=120 -nanopore read_subsets/sample_"$i".fastq
    canu_trim.py canu_temp/canu.contigs.fasta > assemblies/assembly_"$i".fasta
    rm -rf canu_temp
done
for i in 02 08 14 20; do
    flye --nano-hq read_subsets/sample_"$i".fastq --threads 32 --out-dir flye_temp
    cp flye_temp/assembly.fasta assemblies/assembly_"$i".fasta
    cp flye_temp/assembly_graph.gfa assemblies/assembly_"$i".gfa
    rm -r flye_temp
done
for i in 03 09 15 21; do
    miniasm_and_minipolish.sh read_subsets/sample_"$i".fastq 32 > assemblies/assembly_"$i".gfa
    any2fasta assemblies/assembly_"$i".gfa > assemblies/assembly_"$i".fasta
done
for i in 04 10 16 22; do
    ~/programs/NECAT/Linux-amd64/bin/necat.pl config config.txt
    realpath read_subsets/sample_"$i".fastq > read_list.txt
    sed -i "s/PROJECT=/PROJECT=necat/" config.txt
    sed -i "s/ONT_READ_LIST=/ONT_READ_LIST=read_list.txt/" config.txt
    sed -i "s/GENOME_SIZE=/GENOME_SIZE="$genome_size"/" config.txt
    sed -i "s/THREADS=4/THREADS=32/" config.txt
    ~/programs/NECAT/Linux-amd64/bin/necat.pl bridge config.txt
    cp necat/6-bridge_contigs/polished_contigs.fasta assemblies/assembly_"$i".fasta
    rm -r necat config.txt read_list.txt
done
for i in 05 11 17 23; do
    echo read_subsets/sample_"$i".fastq > input.fofn
    cp ~/programs/NextDenovo/doc/run.cfg nextdenovo_run.cfg
    sed -i "s/genome_size = 1g/genome_size = "$genome_size"/" nextdenovo_run.cfg
    sed -i "s/parallel_jobs = 20/parallel_jobs = 1/" nextdenovo_run.cfg
    sed -i "s/read_type = clr/read_type = ont/" nextdenovo_run.cfg
    sed -i "s/pa_correction = 3/pa_correction = 1/" nextdenovo_run.cfg
    sed -i "s/correction_options = -p 15/correction_options = -p 32/" nextdenovo_run.cfg
    sed -i "s/-t 8/-t 32/" nextdenovo_run.cfg
    nextDenovo nextdenovo_run.cfg
    cp 01_rundir/03.ctg_graph/nd.asm.fasta nextdenovo_temp.fasta
    rm -r 01_rundir nextdenovo_run.cfg input.fofn
    echo read_subsets/sample_"$i".fastq > lgs.fofn
    cat ~/programs/NextPolish/doc/run.cfg | grep -v "sgs" | grep -v "hifi" > nextpolish_run.cfg
    sed -i "s/parallel_jobs = 6/parallel_jobs = 1/" nextpolish_run.cfg
    sed -i "s/multithread_jobs = 5/multithread_jobs = 32/" nextpolish_run.cfg
    sed -i "s|genome = ./raw.genome.fasta|genome = nextdenovo_temp.fasta|" nextpolish_run.cfg
    sed -i "s|-x map-ont|-x map-ont -t 32|" nextpolish_run.cfg
    nextPolish nextpolish_run.cfg
    cp 01_rundir/genome.nextpolish.fasta assemblies/assembly_"$i".fasta
    rm -r 01_rundir pid*.log.info nextpolish_run.cfg lgs.fofn nextdenovo_temp.fasta
done
for i in 06 12 18 24; do
    raven --threads 32 --disable-checkpoints --graphical-fragment-assembly assemblies/assembly_"$i".gfa read_subsets/sample_"$i".fastq > assemblies/assembly_"$i".fasta
done
```

```bash
cd /home/wickr/operation_crayfish/plasmids
trycycler cluster --assemblies assemblies/*.fasta --reads plasmid_filtlong.fastq.gz --out_dir trycycler --threads 48
trycycler reconcile --reads plasmid_filtlong.fastq.gz --cluster_dir trycycler/cluster_001
trycycler reconcile --reads plasmid_filtlong.fastq.gz --cluster_dir trycycler/cluster_002
trycycler msa --threads 32 --cluster_dir trycycler/cluster_001
trycycler msa --threads 32 --cluster_dir trycycler/cluster_002
trycycler partition --reads plasmid_filtlong.fastq.gz --cluster_dirs trycycler/cluster_* --threads 48
trycycler consensus --cluster_dir trycycler/cluster_001
trycycler consensus --cluster_dir trycycler/cluster_002
```

```bash
cd /home/wickr/operation_crayfish/plasmids
for c in trycycler/cluster_*; do
    medaka_consensus -i "$c"/4_reads.fastq -d "$c"/7_final_consensus.fasta -o "$c"/medaka -m r1041_e82_400bps_sup_v4.3.0 -t 12
    mv "$c"/medaka/consensus.fasta "$c"/8_medaka.fasta
    rm -r "$c"/medaka "$c"/*.fai "$c"/*.mmi  # clean up
done
```


# Inspect Medaka changes

Medaka didn't make any changes to the plasmids and only made one change to the chromosome:
```
  before medaka 740582-740612: GATGTTGGTAGTATCGGGGGGGGGGGGGGGC
  after medaka  740582-740611: GATGTTGGTAGTATC-GGGGGGGGGGGGGGC
```

Since I don't have any related genome to use for comparison, I assessed the change using my [`mean_prodigal_length.sh`](https://github.com/rrwick/Perfect-bacterial-genome-tutorial/blob/main/scripts/mean_prodigal_length.sh) script.

The pre-Medaka version (15xG) got a slightly better mean Prodigal length than the post-Medaka version (14xG), so I'll use that one going forward. I.e. I reject the only Medaka change.


# Medaka v2.0.1

I came back to this in Oct 2024 because there is now a new version of Medaka with a bacterial-specific model:
```bash
conda activate medaka

cd /home/wickr/operation_crayfish
for c in chromosome/trycycler/cluster_* plasmids/trycycler/cluster_*; do
    medaka_consensus -i "$c"/4_reads.fastq -d "$c"/7_final_consensus.fasta -o "$c"/medaka -m r1041_e82_400bps_bacterial_methylation -t 12
    mv "$c"/medaka/consensus.fasta "$c"/9_medaka_v2.fasta
    rm -r "$c"/medaka "$c"/*.fai "$c"/*.mmi  # clean up
done
```

For all 3 replicons, the `9_medaka_v2.fasta` file was identical to the `7_final_consensus.fasta` file, so Medaka v2 made no changes. This also increases my confidence in the long homopolymer - the fact that Medaka v2 didn't change its length supports my decision to use the pre-Medaka sequence.


# Protein clustering - chromosome

This genome is very repetitive - I suspect ISs or something similar.

```bash
cd ~/Dropbox/Uni_research/Projects/2023-12_Operation_Crayfish/Coxiellaceae_genome/chromosome
prodigal -i trycycler.fasta -a trycycler.faa
seqtk seq trycycler.faa > temp.faa && mv temp.faa trycycler.faa
cd-hit -i trycycler.faa -o trycycler_90.faa
```

Get the clusters with 10 or more sequences:
```python
import collections

clusters = collections.defaultdict(list)
rep_seqs = {}
with open('trycycler_90.faa.clstr', 'rt') as f:
    for line in f:
        if line.startswith('>'):
            current_cluster = line.strip()[1:]
        else:
            seq_name = line.split('>')[1].split('.')[0]
            clusters[current_cluster].append(seq_name)
            if '*' in line:
                rep_seqs[current_cluster] = seq_name

big_clusters = []
for cluster, seq_names in clusters.items():
    if len(seq_names) > 10:
        big_clusters.append((cluster, len(seq_names), rep_seqs[cluster]))

big_clusters = sorted(big_clusters, key=lambda x: x[1], reverse=True)

cluster_seqs = {}
with open('trycycler_90.faa', 'rt') as f:
    for header in f:
        seq = next(f).strip()
        seq_name = header[1:].split(' ')[0]
        cluster_seqs[seq_name] = seq

with open('trycycler_90_big_clusters.faa', 'wt') as f:
    for cluster, cluster_len, cluster_rep in big_clusters:
        f.write(f'>{cluster_rep} cluster_size={cluster_len}\n')
        f.write(f'{cluster_seqs[cluster_rep]}\n')
```


# Protein clustering - shared

```bash
cd ~/Dropbox/Uni_research/Projects/2023-12_Operation_Crayfish/Coxiellaceae_genome
prodigal -i trycycler.fasta -a trycycler.faa
seqtk seq trycycler.faa > temp.faa && mv temp.faa trycycler.faa
cd-hit -i trycycler.faa -o trycycler_90.faa
```

Get the clusters shared between the chromosome and plasmids:
```python
import collections

clusters = collections.defaultdict(list)
rep_seqs = {}
with open('trycycler_90.faa.clstr', 'rt') as f:
    for line in f:
        if line.startswith('>'):
            current_cluster = line.strip()[1:]
        else:
            seq_name = line.split('>')[1].split('.')[0]
            clusters[current_cluster].append(seq_name)
            if '*' in line:
                rep_seqs[current_cluster] = seq_name

shared_clusters = []
for cluster, seq_names in clusters.items():
    chromosome_names = [n for n in seq_names if n.startswith('chromosome')]
    plasmid_names = [n for n in seq_names if n.startswith('plasmid')]
    if chromosome_names and plasmid_names:
        shared_clusters.append((cluster, len(seq_names), len(chromosome_names), len(plasmid_names), rep_seqs[cluster]))

shared_clusters = sorted(shared_clusters, key=lambda x: x[1], reverse=True)

cluster_seqs = {}
with open('trycycler_90.faa', 'rt') as f:
    for header in f:
        seq = next(f).strip()
        seq_name = header[1:].split(' ')[0]
        cluster_seqs[seq_name] = seq

with open('trycycler_90_shared_clusters.faa', 'wt') as f:
    for cluster, cluster_len, chromosome_count, plasmid_count, cluster_rep in shared_clusters:
        cluster_num=cluster.replace('Cluster ', '')
        f.write(f'>{cluster_rep} cluster={cluster_num} cluster_size={cluster_len} chromosome_count={chromosome_count} plasmid_count={plasmid_count}\n')
        f.write(f'{cluster_seqs[cluster_rep]}\n')
```


# Genome completeness

I'm reasonably confident that my assembled chromosome (2233862 bp) and two plasmids (59783 bp and 34021 bp) all came from the same genome. This is because:
* They all have similar depths in the assemblies, e.g. 2338x, 1834x and 1865x in the Flye assembly.
* They have a lot of shared genes. 17 of the cd-hit clusters have genes in the chromosome and at least one of the plasmids.
* When I tblastn (70% coverage and 90% identity thresholds) for those 17 clusters in my full metagenome assemblies, I only get hits in contigs from those three replicons.

I'm also reasonably confident that I didn't miss any replicons from that genome, because when I filter for high-depth contigs (e.g. 1000x or more), I only find:
* those three replicons
* stuff tangled into the crayfish part of the graph - presumably repetitive elements
* extremely repetitive sequences, e.g. `CCATT` thousands of times (ONT junk?)


# Rotation

While Trycycler didn't rotate to a dnaA gene (not close enough identity), I manually searched and found this gene:
```
GTGTCATTGTCAACAAACTTATGGGAGAAGTGTCTCAATTATTTAAGAGACGAAATCCCGCCACAACAATATAATACGTGGATTCGACCATTACATGTTGTAGAACAACATAATGGTTTATTATTGCTCGCGCCCAATCGATTTGTGCTCGATTGGATCAACGAGCGGTTTTTAACCCGTATCAATGAATTACTGGCAGAATTGTCCGAAGACGCCCCACCTCGAATTAAATTACAAATTGGCAGTAAAACAACTAACGAAGATAATAACACTCCCAACGTCTCTAAACCAAATAGTCCAGCCGTTCACACACAAACATTTAGTAACACGAATGGTAAGTTACAGTTTGGTACGATGAACCCGCATGGTAATTTGAATTCTCATTTTACGTTTGAGAACTTTGTAGAAGGTAAATCAAATCAATTAGCACGTGCAGCCGCCACACAAGTCGCGAAAAATCCAGGGCAAGCTTATAACCCGCTCTTTATATATGGTGGTGTAGGCCTCGGTAAAACACATTTAATGCACGGCATCGGAAATTTTATTTTACAGAACAACCCCAATAGAAAAGTTTTATATCTACACTCCGAACGATTTGTTGCGGATATGATTAAAGCTTTACAACGAAATGCTATGAATGAATTTAAACGTTATTATCGTTCGGTAAACGCACTCTTAATTGACGACATTCAATTTTTTGCAGGTAAAGATCGTTCTCAAGAAGAATTTTTTCACACATTTAATGCATTATTGGATGGGCAGCAACAAATAATTCTCACGTGCGATCGCTATCCCAAAGAAATTGACGGCCTTGAAGAGCGTTTGCAATCACGTTTCGGGTGGGGCCTGACAGTCGCAATTGAACCCCCCGAACTCGAAACTCGCGTTGCAATTTTAATGACAAAAGCAGAACAGTTAAAAATCGCTCTTCCCCACGAAGTTGCATTTTTCATTGCAAAAAATATTCAATCGAACGTCCGAGAATTAGAGGGTGCACTAAAAAGAGTCATTGCCAATGCGCACTTCACCGGCCAATCCATTACAGTCGATTTCACCCGCGATGCGCTAAAGGATTTATTGACGTTACAAGCTAAACTCGTAACAATTGAGAACATACAGAGAACGGTCGCGGAGTATTATAAAATTAAAGTAGCTGATTTGTTATCAAAACGTCGCAGCCGCTCTATTGCCAGGCCACGACAAATTGCAATGACTCTCTCAAAAGAGCTGACTAATCATAGTCTTCCTGAAATTGGTGATGCTTTTGGCGGTCGAGATCACACCACCGTGTTGCATGCCTGCCGAAAAATAAAAGAATTACTAGCAAGTTCTTTAGATATACAAGAAGATTATAAAAATTTGTTACGAATTTTGTCCGCTTAA
```
Which is a pretty good match, so I rotated the chromosome to start there.


# Tree

Gather GTDB assemblies in Coxiellaceae (also all of Coxiellales since there is just one family in that order):
```bash
cd ~/operation_crayfish/tree

for g in 2-02-FULL-42-43 2-12-FULL-42-15 CACHNP01 CAIQCE01 Coxiella GCA-2732805 JABABH01 JAFEDN01 JAHFRQ01 JAHZIV01 JAHZKQ01 JAJFJO01 JAJRRL01 UBA9148; do
    tar -xvf /home/db/kraken2/gtdb_r214/assemblies_100_per_species.tar assemblies/"$g"'_*' &
done

for f in assemblies/*/*.fna.gz; do
    new=$(echo "$f" | sed 's|/GC|_GC|')
    mv "$f" "$new"
done
mv assemblies/GCA-2732805_sp002732805/GCA_002732805.1.fna.gz assemblies/GCA-2732805_sp002732805_GCA_002732805.1.fna.gz
```

Dereplicate a bit (mostly culling _Coxiella burnetii_ genomes):
```bash
dereplicator.py --threads 32 --distance 0.001 assemblies derep
rm -r assemblies mash.phylip 
mv derep assemblies
```

I then manually added `crayfish.fna` to the `assemblies` directory and built a tree:
```bash
cd ~/operation_crayfish/tree
gunzip assemblies/*.fna.gz
conda activate gtdbtk
export GTDBTK_DATA_PATH=/home/wickr/release214
gtdbtk de_novo_wf --genome_dir assemblies --bacteria --outgroup_taxon o__Enterobacterales --out_dir gtdbtk_tree --cpus 64

gtdbtk identify --genome_dir assemblies --out_dir gtdbtk_1_identify --cpus 64
gtdbtk align --identify_dir gtdbtk_1_identify --out_dir gtdbtk_2_align --cpus 64
gtdbtk infer --msa_file gtdbtk_2_align/align/gtdbtk.bac120.user_msa.fasta.gz --out_dir gtdbtk_3_infer --cpus 64
```

Minimum-variance rooting:
```bash
FastRoot.py -i gtdbtk.unrooted.tree -o coxiellaceae.newick
```


# Annotation

Using Bakta v1.9.1 with:
* Bakta database 5.0
* AMRFinderPlus database 2023-11-15.1

```bash
cd ~/operation_crayfish
export BAKTA_DB=/home/wickr/bakta_db_5.0
bakta --output bakta --prefix coxiella_cheraxi coxiella_cheraxi_genome.fasta
```

I then manually had to add `/db_xref="taxon:426622"` to the source feature lines, in order for [S4TE](https://sate.cirad.fr/S4TE.php) to accept it.
